# Propagation of Beta/Gamma Rhythms in the Cortico-Basal Ganglia Circuits of the Parkinsonian Rat

**DOI:** 10.1101/180455

**Authors:** Timothy O. West, Luc Berthouze, David M. Halliday, Vladimir Litvak, Andrew Sharott, Peter J. Magill, Simon F. Farmer

## Abstract

Some motor impairments associated with Parkinson’s disease are thought to arise from pathological activity in the neuronal networks formed by the basal ganglia (BG) and motor cortex. To evaluate several hypotheses proposed to explain the emergence of pathological oscillations in Parkinsonism, we investigated changes to the directed connectivity in these networks following dopamine depletion. We recorded local field potentials in the cortex and basal ganglia of anesthetized rats rendered Parkinsonian by injection of 6-hydroxydopamine (6-OHDA), with dopamine-intact rats as controls. We performed systematic analyses of the networks using a novel tool for estimation of directed neural interactions, as well as a conditioned variant which permits the analysis of the dependence of a connection upon a third reference signal. We find evidence of the dopamine dependency of both low beta (14-20 Hz) and high beta/low gamma (20-40 Hz) directed interactions within the BG and cortico-BG networks. Notably, 6-OHDA lesions were associated with enhancement of the cortical “hyperdirect” connection to the subthalamic nucleus (STN), as well the STN’s feedback to the cortex and striatum. We find beta synchronization to be robust to conditioning using signals from any one structure. Conversely, we find that high beta/gamma drive from the cortex to subcortical regions is weakened by 6-OHDA lesions and is susceptible to conditioning. Furthermore, we provide evidence that gamma is routed from striatum in a pathway that is independent of STN. These results further inform our understanding of the substrates for pathological rhythms in salient brain networks in Parkinsonism.

**New & Noteworthy:** We present a novel analysis of electrophysiological recordings in the cortico-basal ganglia network with the aims of evaluating several hypotheses concerning the origins of abnormal brain rhythms associated with Parkinson’s disease. We present evidence for changes in the directed connections within the network following chronic dopamine depletion in rodents. These findings speak to the plausibility of a “short-circuiting” of the network that gives rise to the conditions from which pathological synchronization may arise.

## Introduction

The basal ganglia (BG) are host to a small but important cluster of dopaminergic neurons that act to modulate the activity of a large re-entrant network that comprises the cortico-basal ganglia-thalamo-cortical circuit (DeLong and Wichmann 2010; Lanciego et al. 2012). Investigation of the structure of this network (Smith et al. 1998; Bolam et al. 2000) has led to a canonical view of the circuit (depicted in figure 1) from which multiple process theories of BG function have arisen (for a review, see Schroll and Hamker (2013)).

**Figure 1.**
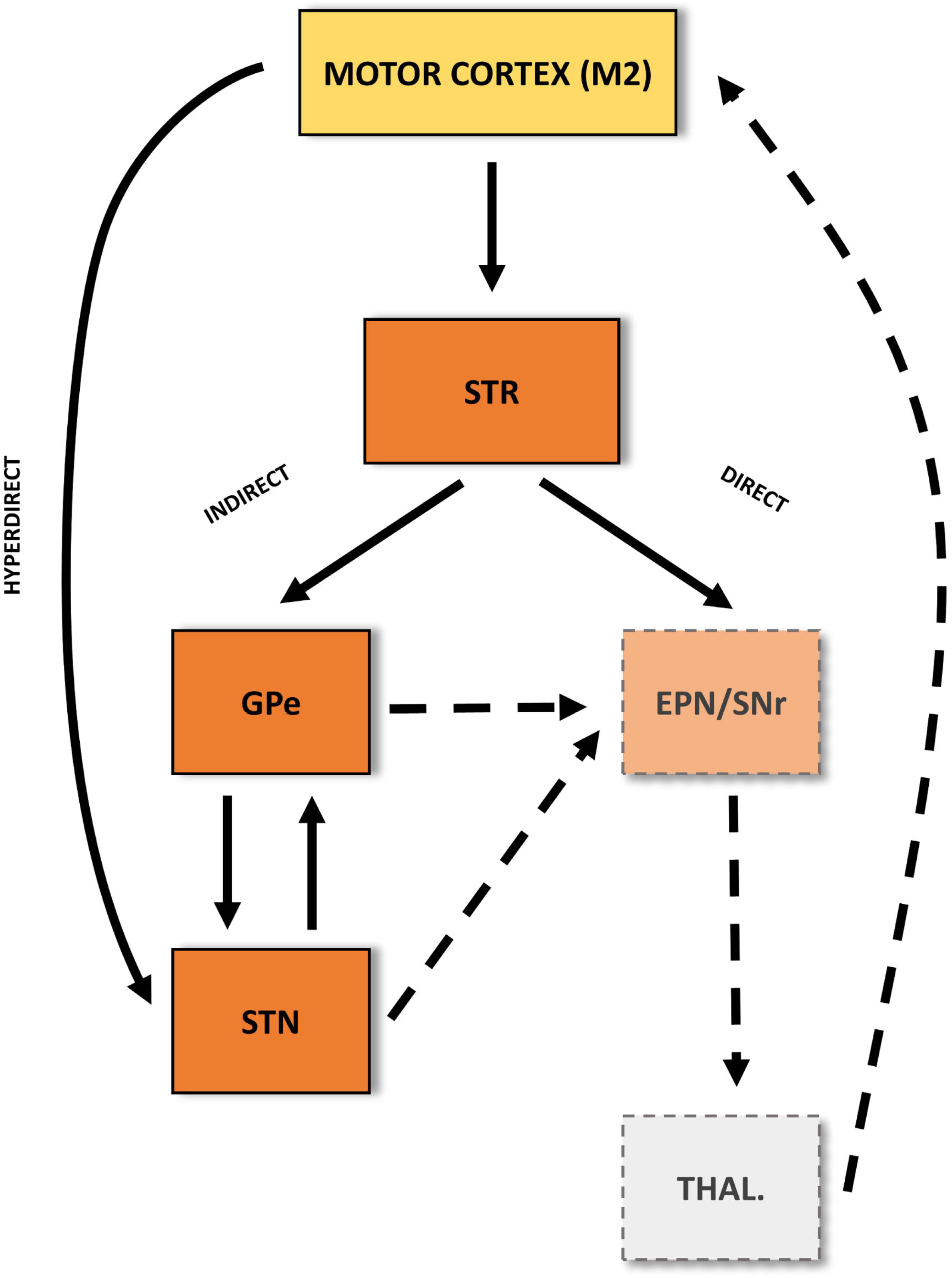
Schematic of canonical cortical-basal ganglia circuit incorporating the direct and indirect pathways, as well as the cortico-subthalamic hyperdirect pathway. The motor cortex (M2, yellow box) has major inputs to the basal ganglia (BG, orange boxes) at the striatum (STR) and subthalamic nucleus (STN). Information flow along the indirect, direct and hyperdirect pathways ultimately impinges on the output nuclei of the basal ganglia, made up of the entopeduncular nucleus (EPN) and substantia nigra pars reticulata (SNr). BG output targets thalamic relays, of which some return back to M1. Structures for which experimental data is available are shown in bold outline, otherwise in dashed.

Recent theory concerning the organisation of brain networks and communication within them via synchronized oscillations (Varela et al. 2001; Fries 2005, 2015; Bressler and Menon 2010; Thut et al. 2012) has emphasised the importance of understanding the dynamics of these networks beyond that afforded by studying structure alone (Deco et al. 2008, 2012). Neural oscillations and their synchronization can be detected across multiple scales of brain activity, from single neuronal discharges up to the level of mesoscale neural ensembles such as those measured in the local field potential (LFP) or electrocorticogram (ECoG). Pathological oscillations and their synchronization across disparate regions of the brain have been detected in brain disorders such as Parkinson’s disease (PD), schizophrenia, and epilepsy, leading to the hypothesis that the oscillations themselves bear a causal role in their associated behavioral impairments (Schnitzler and Gross 2005; Uhlhaas and Singer 2006; Hammond et al. 2007).

Excessive beta oscillations (14-30 Hz) in the BG associated with dopamine depletion have been observed reliably in untreated patients with PD (Levy et al. 2000; Brown et al. 2001; Weinberger et al. 2006; Hammond et al. 2007). Beta rhythms are attenuated by treatments such as dopamine replacement therapy (Kühn et al. 2006; Weinberger et al. 2006; West et al. 2016; Beudel et al. 2017; Levy 2002) and deep brain stimulation (DBS) (Ray et al. 2008; Eusebio et al. 2011; Whitmer et al. 2012) in a way that correlates with the degree of the improvement of motor symptoms. This has strengthened the argument that the pathological beta rhythms are directly related to the functional impairment seen in patients (Hanslmayr et al. 2012; Brittain and Brown 2014). Furthermore, gamma activity in the motor system has been hypothesized to be prokinetic (Schoffelen et al. 2005). In PD, the spectral power of multiunit recordings from STN at 40-90 Hz have been demonstrated to be negatively correlated with bradykinetic symptoms in patients (Sharott et al. 2014).

The pathological oscillations observed in mesoscale electrophysiological signals are a direct consequence of changes to the underlying networks of neuronal ensembles that generate them. This understanding has led to the re-classification of multiple neurological diseases such as PD or Tourette’s as ‘circuit disorders’ (DeLong and Wichmann 2010). Knowledge of how dopamine depletion results in changes to the network, and the subsequent emergence of pathological synchrony is likely to lead to a better understanding of the causes of impairment and its treatments (Shen et al. 2008; Schroll et al. 2014). Thus improving insight into how changes to connections in the network lead to the emergence of pathological dynamics remains an important line of enquiry (Wichmann and DeLong 1999; Dostrovsky and Bergman 2004; Holgado et al. 2010)

Previous work aiming to understand the origins of the pathological beta rhythm has involved systematic lesioning of the BG network (Ni et al. 2000; Tachibana et al. 2011), computational modelling (Holgado et al. 2010; Moran et al. 2011; Marreiros et al. 2013; Nevado-Holgado et al. 2014; Pavlides et al. 2015), and techniques from signal analysis (Sharott et al. 2005a; Mallet et al. 2008a, 2008b; Litvak et al. 2011). In this paper, we take the latter approach and, through analysis of neural recordings, aim to infer the changes in neural transmission that occur in cortico-BG circuits following chronic dopamine depletion.

Connectivity between parts of the brain can be inferred from the statistical dependencies that arise due to neural transmission: we refer to this as functional connectivity (Friston 2011). Previous studies have aimed to describe ‘effective’ connectivity (i.e. causal interactions) within this network and have employed the dynamic causal modeling (DCM) framework in order to do so. To date, two such studies have utilised the inversion of biophysical models upon cross spectral densities from recordings in either anaesthetised 6-OHDA lesioned rats (Moran et al. 2011), or awake DBS patients (Marreiros et al. 2013). Both found evidence for the strengthening of the cortico-subthalamic connection (termed the ‘hyper-direct’ pathway (Nambu et al. 2002)) in the dopamine-depleted state.

From this work amongst others, several hypotheses have arisen concerning the emergence of pathological beta rhythms as a result of the dopamine depletion associated with PD (for a review see Holgado et al. 2010). These include the dopamine-dependent modulation of recurrent loops within the network, either between the reciprocally-coupled network of neurons of the subthalamic nucleus (STN) and the external globus pallidus (GPe) (Plenz and Kital 1999; Bevan et al. 2002; Terman et al. 2002; Holgado et al. 2010); or of a longer loop involving feedback from BG output nuclei to the cortex via thalamo-cortical tracts (Leblois et al. 2006; Pavlides et al. 2012, 2015). Alternatively, it has been proposed that dopamine depletion disrupts mechanisms which regulate the gain of cortical afferents to the BG and somehow disrupt striatal outflow (Brown 2007; Hammond et al. 2007).

Here, using a recently described non-parametric (model-free) signal analysis technique (Halliday et al., 2015), we study the effects of dopamine depletion upon neural connectivity in the network formed by elements of the BG and motor cortex in 6-OHDA-lesioned and dopamine-intact control rats. We employ this method as a measure of *directed* functional connectivity (hereon shortened to directed connectivity). It is a model-free estimate that makes no assumptions as to the causes of the data (Bastos and Schoffelen 2016), only that temporal precedence implies a driving neuronal influence (please see later sections for discussion). Furthermore, we use a a multivariate extension of the framework (Halliday et al. 2016) in order to determine whether the interaction between two areas shares correlation with activity recorded at a third structure in the network. This approach provides insight into frequency-specific directional connectivity and the dependence of coupled signals upon activity at a third region. By recording LFPs and ECoG in 6-OHDA–lesioned animals and dopamine-intact controls we aim to identify changes to connectivity that occur as a result of the loss of dopamine from these circuits. Our findings are interpreted within the context of the canonical circuit (figure 1), other existing models of basal ganglia connectivity, and several hypotheses concerning the generation and propagation of pathological beta rhythms in the network.

## Methods

### Experimental Data

#### Electrophysiological Recordings

Experimental procedures were carried out on adult male Sprague-Dawley rats (Charles River, Margate, UK) and were conducted in accordance with the Animals (Scientific Procedures) Act, 1986 (UK). Recordings were made in eight dopamine-intact control rats (288–412 g) and nine 6-OHDA-lesioned rats (285– 428 g at the time of recording), as described previously (Magill et al. 2006; Mallet et al. 2008a, 2008b; Moran et al. 2011). Briefly, anaesthesia was induced with 4% v/v isoflurane (Isoflo, Schering-Plough Ltd., Welwyn Garden City, UK) in O2, and maintained with urethane (1.3 g/kg, i.p.; ethyl carbamate, Sigma, Poole, UK), and supplemental doses of ketamine (30 mg/kg; Ketaset, Willows Francis, Crawley, UK) and xylazine (3 mg/kg; Rompun, Bayer, Germany).

The ECoG was recorded via a 1 mm diameter steel screw juxtaposed to the dura mater above the right frontal (somatic sensory-motor) cortex (4.5 mm anterior and 2.0 mm lateral of Bregma and approximately centred over M2, (Paxinos and Watson 2007)) and was referenced against another screw implanted in the skull above the ipsilateral cerebellar hemisphere. Raw ECoG was band-pass filtered (0.3–1500 Hz, −3 dB limits) and amplified (2000x; DPA-2FS filter/amplifier: Scientifica Ltd., Harpenden, UK) before acquisition. Extracellular recordings of LFPs in the dorsal striatum (STR), GPe and STN were simultaneously made in each animal using ‘silicon probes’ (NeuroNexus Technologies, Ann Arbor, MI). Each probe had one or two vertical arrays of recording contacts (impedance of 0.9–1.3 MΩ measured at 1000 Hz; area of ~400μm^2^).

The same probe was used throughout these experiments but it was cleaned after each experiment in a proteolytic enzyme solution to ensure that contact impedances and recording performance were not altered by probe use and re-use. Monopolar probe signals were recorded using high-impedance unity-gain operational amplifiers (Advanced LinCMOS: Texas Instruments, Dallas, TX) and were referenced against a screw implanted above the contralateral cerebellar hemisphere. After initial amplification, extracellular signals were further amplified (1000x) and low-pass filtered at 6000 Hz using programmable differential amplifiers (Lynx-8: Neuralynx, Tucson, AZ). The ECoG and probe signals were each sampled at 17.9 kHz using a Power1401 Analog-Digital converter and a PC running Spike2 acquisition and analysis software (Cambridge Electronic Design Ltd., Cambridge, UK). Recording sites in the BG were verified by post hoc histology, as described previously (Magill et al. 2006; Mallet et al. 2008a, 2008b).

Neuronal activity was recorded during episodes of spontaneous ‘cortical activation’, which contain patterns of activity that are similar to those observed during the awake, behaving state (Steriade 2000). Cortical activation was defined according to ECoG activity. Neuronal activity patterns present under this anaesthetic regime may only be qualitatively similar to those present in the un-anesthetized brain. However, the urethane-anesthetized animal still serves as a useful model for assessing ensemble dynamics within the basal ganglia. Indeed, in 6-OHDA-lesioned animals, exaggerated beta oscillations emerge in cortico-basal ganglia circuits during activated brain states thus accurately mimicking the oscillatory activity recorded in awake, un-medicated PD patients. Examples of the raw and pre-processed electrophysiological signals as well the corresponding power spectra for control and lesioned animals are shown in figure 2.

**Figure 2.**
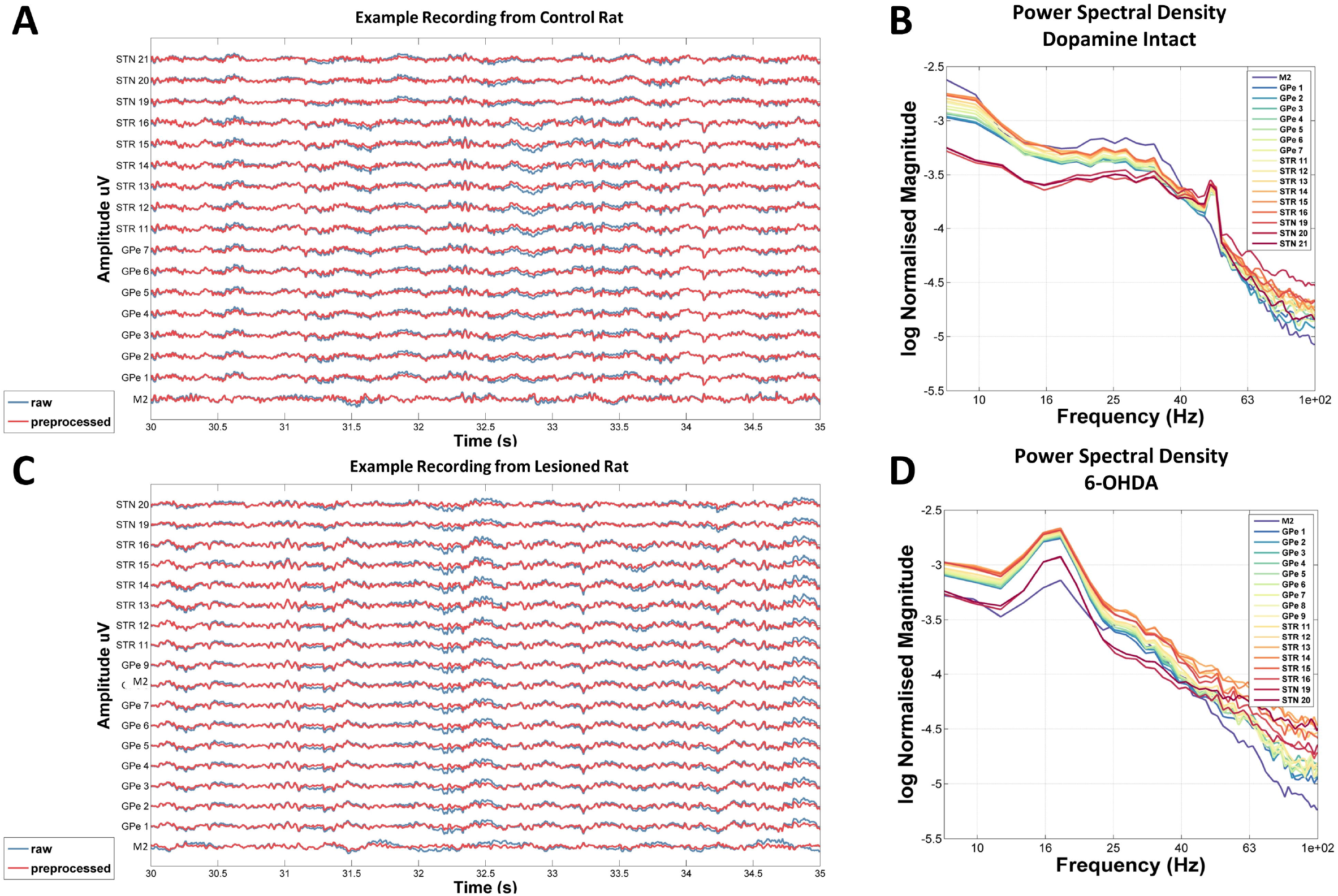
Example outcome of data pre-processing of rat subcortical monopolar LFP and cortical ECoG signals for a single animal from the control (A-B) or Parkinsonian (C-D) groups. (A) 5 second excerpt of recording from one dopamine intact, control animal. The example trace shows the time course of LFP recordings recorded using silicon electrodes implanted in the external globus pallidus (GPe), striatum (STR) and subthalamic nucleus (STN), and with multiple contacts within each structure. Additionally, there is a cortical ECoG signal recorded from a screw over motor cortex (M2). Raw data is shown in blue whilst pre-processed signals are overlaid in red. The data was demeaned and then high pass filtered at 4 Hz. (B) Spectral analysis of example control animal’s recording. Data epoched into 2-second segments contaminated by high amplitude transients were removed using Z-thresholding as described in the text. (C) Same as (A) but for an example 6-OHDA-lesioned (dopamine depleted) animal. (D) Same as (B) but for 6-OHDA-lesioned animal.

#### 6-Hydroxydopamine Lesions of Dopamine Neurons

Unilateral 6-OHDA lesions were carried out on 200–250 g rats, as described previously (Mallet et al. 2008a, 2008b). Twenty-five minutes before the injection of 6-OHDA, all animals received a bolus of desipramine (25 mg/kg, i.p.; Sigma) to minimize the uptake of 6-OHDA by noradrenergic neurons (Schwarting and Huston 1996a). Anaesthesia was induced and maintained with 4% v/v isoflurane (see above). The neurotoxin 6-OHDA (hydrochloride salt; Sigma) was dissolved immediately before use in ice-cold 0.9% w/v NaCl solution containing 0.02% w/v ascorbate to a final concentration of 4 mg/ml. Then 3 ml of 6-OHDA solution was injected into the region adjacent to the medial substantia nigra (4.5 mm posterior and 1.2 mm lateral of bregma, and 7.9 mm ventral to the dura. The extent of the dopamine lesion was assessed 14–16 days after 6-OHDA injection by challenge with apomorphine (0.05 mg/kg, s.c.; Sigma) (Schwarting and Huston 1996b). The lesion was considered successful in those animals that made >80 net contraversive rotations in 20 min. Electrophysiological recordings were carried out ipsilateral to 6-OHDA lesions in anesthetized rats 21–42 days after surgery, when pathophysiological changes in the basal ganglia are likely to have levelled out near their maxima (Mallet et al. 2008a).

### Data Acquisition and Analysis

#### Data Conversion and Pre-Processing

To isolate LFPs and ECoGs, all electrophysiological data were down-sampled from a hardware native 17.9 kHz to 250 Hz using Spike2 acquisition and analysis software (version 4; Cambridge Electronic Design Ltd., Cambridge, UK). Data were then imported from Spike2 into MATLAB (The Mathworks, Nantucket, MA, USA) where they were analysed using custom scripts utilizing routines from the Fieldtrip software package (packaged in SPM 12.3) (Oostenveld et al. 2011), as well as Neurospec (http://www.neurospec.org/). Data were pre-processed as follows: i) data were first truncated to remove 1 second from either end of the recording, ii) mean subtracted; iii) band-passed filtered with a finite impulse response, two-pass (zero-lag) filter designed such that the filter order is rounded to the number of samples for 3 periods of the lowest frequency, between 4-100 Hz; iv) data were then split into 1 second epochs; v) each epoch was subjected to a Z-score threshold criterion such that epochs containing any high amplitude artefacts were removed. Examples of outcomes from this pre-processing are shown in figure 2.

#### Signal Analysis Techniques

Power analyses were made using the averaged periodogram method across 1 second epochs and using a Hanning taper to reduce the effects of spectral leakage. Frequencies between 49-51 Hz were removed so that there was no contribution from 50 Hz line noise. We computed spectral power, imaginary coherence, directional analyses (NPD), and their conditioned variants using the Neurospec analysis toolbox.

##### Non-zero Phase Lag Functional Connectivity Analysis: Imaginary Coherence

The standardly used spectral coherence (Halliday et al. 1995) is sensitive to spurious correlations resulting from instantaneous volume conduction between the two signals of interest (Bastos and Schoffelen 2016). In order to circumvent this issue, several methods have been developed such as taking the imaginary part of coherence (Nolte et al. 2004), the phase lag index (PLI) (Stam et al. 2007), or the weighted phase lag index (Vinck et al. 2011). For this study, we use arguably the simplest method available (imaginary coherence-iCOH) that is derived from the complex coherency. This measure shares the property of disregarding zero lag correlations with the non-parametric directionality analysis that we later introduce for estimates of directed connectivity. We also note the concerns in Stam et al. (2007) on the validity of imaginary coherence analysis, so include additional analyses based on non-parametric directionality.

##### Non-Parametric Directionality

Estimates of directed connectivity were computed using non-parametric directionality (NPD) (Halliday 2015) as implemented in the Neurospec software package. This analysis combines Minimum Mean Square Error (MMSE) pre-whitening with forward and reverse Fourier transforms to decompose coherence estimates at each frequency into three components: forward, reverse and zero lag. These components are defined according to the corresponding time lags in the cross-correlation function derived from the MMSE pre-whitened cross-spectrum. The advantage of this approach is that it allows us to decompose the signal into distinct forward and reverse components of coherence separate from the zero-lag (or instantaneous) component of coherence which is assumed to reflect volume conduction. The method uses temporal precedence to determine directionality. For example, STN activity lagging M2 activity results in a significant forward component of coherence between M2 and STN (when M2 is assumed to be the reference), whereas STN activity leading M2 activity results in a significant reverse component of coherence.

In addition we used a multivariate extension of the non-parametric directionality (Halliday et al. 2016), which allows the directional components of coherence to be conditioned on a third signal. This analysis decomposes the partial coherence into the same three directional components: forward, reverse and zero-lag. This analysis can indicate if the signals reflected in the correlation are common to other parts of the network. For example, the partial correlation between A and B with C as predictor can be used to determine if the flow of information from A → B is independent of area C, or whether the flow of information is A → C → B, in which case the partial coherence between A and B with C as predictor should be zero. The partial coherence can also be used to investigate if the flow of information is C → A and C → B, or if it is A → B → C or C → A → B, which for the latter case the partial coherence, and any directional components should be zero.

This assumes that the conditioning signal, C, is representative of the activity in the relevant brain area. If the signal, C, only captures part of the activity in the brain area then the partial coherence estimate may still have residual features. The most robust interpretation of the partial coherence and multivariate non-parametric directionality is where the partial coherence (and any directional components) are not significant compared to the directional components for the ordinary coherence.

#### Statistics

In order to make statistical comparisons of power, connectivity and directionality spectra between lesioned and control animals we used cluster based permutation testing (Maris and Oostenveld 2007) which avoids introducing bias through the prior specification of frequency bands. It requires no assumption of normality, and affords a correction for the multiple comparison problem by controlling the family-wise error rate via an approximation of the reference distribution for the test statistic. For more details see Maris (2012). The cluster-forming threshold was p<0.05 and the permutation test threshold was set at p<0.025 (as it is a two-sided test). The number of permutations was set to 5000 which tenders a lowest possible P-value equal to 0.0004. Cluster statistics were computed using the ‘*ft_freqstatistics*’ routine in Fieldtrip.

## Results

### Spectral Power

Examples of spectra computed from LFP and ECoG signals recorded in individual animals can be seen in figure 2 (B and D). All the 6-OHDA-lesioned rats demonstrated a clear peak in the spectra in the range 18-22 Hz (encompassing low beta/lower end of high beta frequencies) for LFP recordings across all subcortical recording sites as well as for the sensorimotor ECoGs. In some animals, cortical beta was weaker than that observed subcortically. None of the LFP data from control animals contained a similar beta peak in the spectra although some (4 of 8) showed a wide, low amplitude peak around 20-40 Hz most prominent in the recordings at M2 (an example of which is seen in figure 2B). Analysis of the group averaged spectra (figure 3) shows that the beta peak is significantly increased in the dopamine-depleted animals. Cluster-based permutation testing demonstrated significant differences in group level spectra between control and lesion conditions with clusters showing increases in power in the STN (16-21 Hz, P=0.023), GPe (17-22 Hz, P=0.009) and STR (18-21 Hz, P=0.019) for the dopamine-depleted animals. Cluster-based permutation did not reveal significant increases in beta power in the ECoG signals recorded in the M2 of lesioned rats when compared to the controls (although a positive cluster was found that exceeded the cluster forming threshold but was not significant under permuation testing). No differences between lesioned and control animals were found for frequencies >22 Hz in any structures.

**Figure 3.**
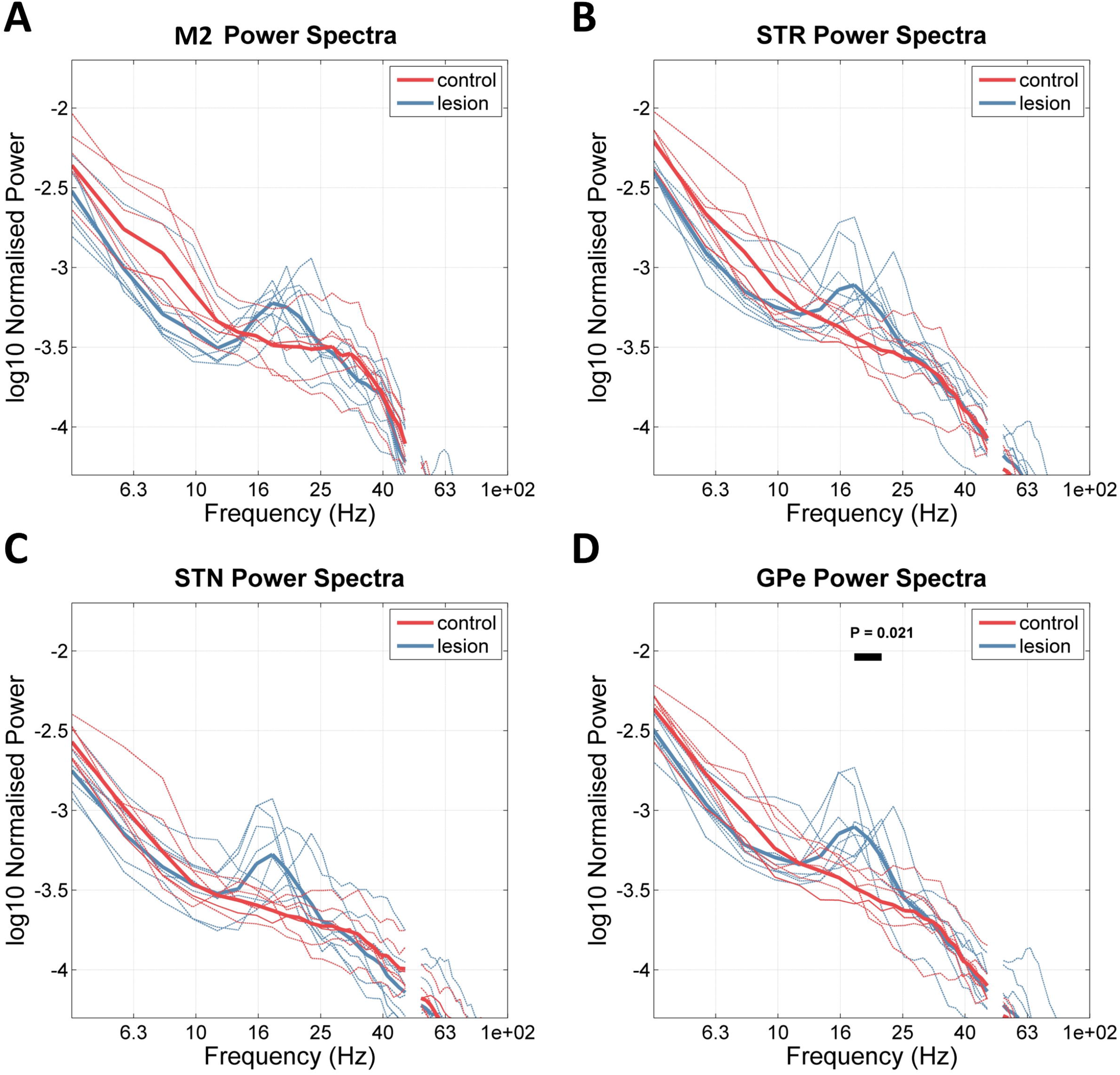
Group averaged power spectra for all rats across both control and lesion conditions. Spectra are shown for signals recorded from (A) motor cortex (M2), (B) the striatum (STR), (C) the subthalamic nucleus (STN), and (D) the external globus pallidus (GPe). Individual spectra are shown in dashed lines whilst the group averages for either the 6-OHDA dopamine depleted or control animals are shown by bold lines in red or blue respectively. Results of cluster permutation tests are indicated by the bar and corresponding P-value. All recording sites presented beta peaks around 18-20 Hz. Cluster based permutation testing for significant differences between conditions showed that there was a significant increase in beta in the lesioned animals for signals recorded at STR, STN, and GPe.

### Functional Connectivity: Imaginary Coherence (iCOH)

Initial analyses of connectivity of the recorded LFPs using magnitude squared coherence showed large magnitude (>0.9) wideband (0 – 60Hz) coherences that were indicative of a large field spread effect (data not shown). This was most apparent in subcortical-subcortical analyses but was also detected for cortical-subcortical pairings. To estimate coherence avoiding contamination by volume conduction we opted to calculate non-zero phase lag correlations using the imaginary part of coherence (iCOH) (see figure 4).

**Figure 4.**
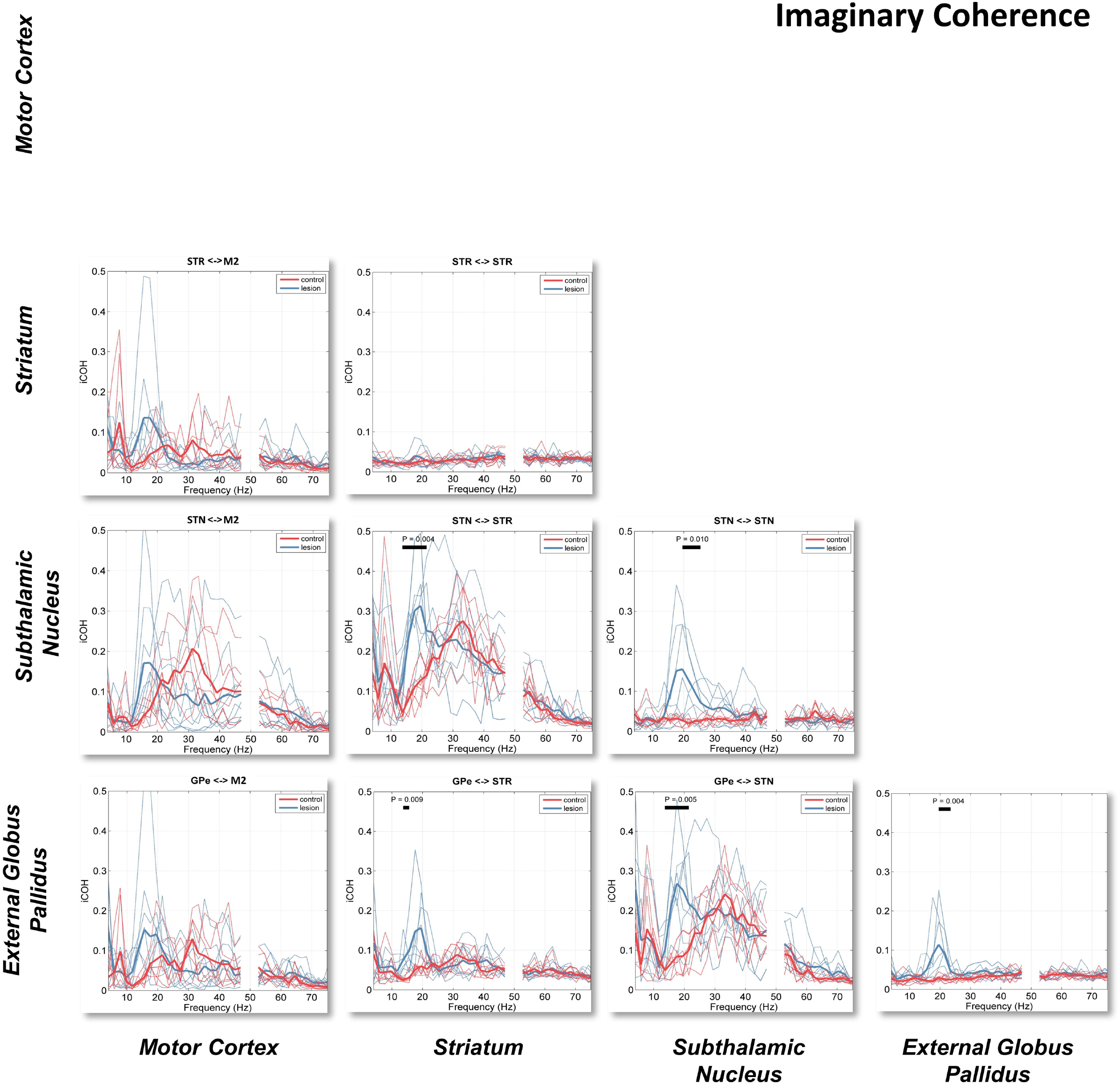
Functional connectivity estimates using imaginary part of coherence (iCOH). Spectra for each animal are shown by thin lines corresponding to either 6-OHDA lesioned (blue) or control (red). The group averages are shown by the bold lines. Cluster-based permutation statistics were applied to determine significant differences between conditions and significant clusters are indicated by the bold line above the spectra and corresponding P-value. The iCOH metric, robust to zero-lag interactions, presents a rich view of functional connectivity that would otherwise be missed if using standard coherence (data not shown). Beta activity is predominant across all cross-regional pairings. STN and GPe also show some intra-nuclear correlations in this range in the dopamine depleted state. Notably there is also a high beta/gamma interaction between STN/M2 and STN/STR that is visible in both control and lesion animals.

We found that activity in the low beta range (14-20 Hz) associated with 6-OHDA dopamine depletion is spread diffusely across the network with all inter-regional comparisons showing a significant beta peak in the iCOH spectrum. Notably, the strongest beta band involved STN, with STN/STR and STN/GPe pairs both showing coefficients greater than 0.2. Within region connectivity (i.e. STN contact 1 to contact 2) was found to be present in this range for only recordings within STN or GPe, where there is a clear beta peak.

Analysis of statistical differences using the cluster based permutation testing between control and lesioned animals showed significant increases of iCOH in the beta band in the lesioned animals and for 5/10 pairs tested: STN/STR (14-21 Hz, P=0.004), STN/STN (19-25 Hz, P = 0.010), GPe/STR (14-16 Hz, P = 0.009), GPe/STN (14-21 Hz, P=0.005), and GPe/GPe (19-23 Hz, P=0.004). Notably, no pairs involving M2 showed significant modulations of beta-band activity with dopaminergic state as detectable with the cluster statistics. Taken generally, these results are indicative of widespread, nonzero lag, low beta-band connectivity across the entire cortico-BG network that is increased in the dopamine-depleted rats.

In the control rats, connectivity in the beta range was reduced. Instead, there was wide-band iCOH in the high beta/low gamma bands, ranging from 20 Hz to 50 Hz in most cases but up to 70 Hz for the STN/M2 interactions. The majority of gamma interactions where iCOH was high (> 0.2) were found in connections involving the STN. Additionally, iCOH in these bands is evident between GPe/M2 and GPe/STR although this was weaker (at around ~0.1) than connections analysed with pairs involving the STN. iCOH in these bands is present in both the lesioned and control animals and does not show a strong modulation by dopamine as evidenced by the lack of significant clusters in the permutation tests for these bands.

### Non-parametric Directionality (NPD)

We next investigated directed connectivity between recorded regions. The results of the analysis using the NPD measure are presented in figure 5. The iCOH and the sum of the non-instantaneous parts (forward and backward) of the NPD are similar, and both methods reveal similar patterns of connectivity (data not shown). Analysis of the instantaneous NPD in isolation demonstrated the existence of high amplitude, wide-band interactions that were similar to those found with magnitude squared coherence (data not shown), and are likely due to zero-phase field spread of activity between recordings. Analyses of directional interactions of the LFPs and ECoG hereon will use the forward and backward components of the NPD to discern directional connectivity between structures.

**Figure 5.**
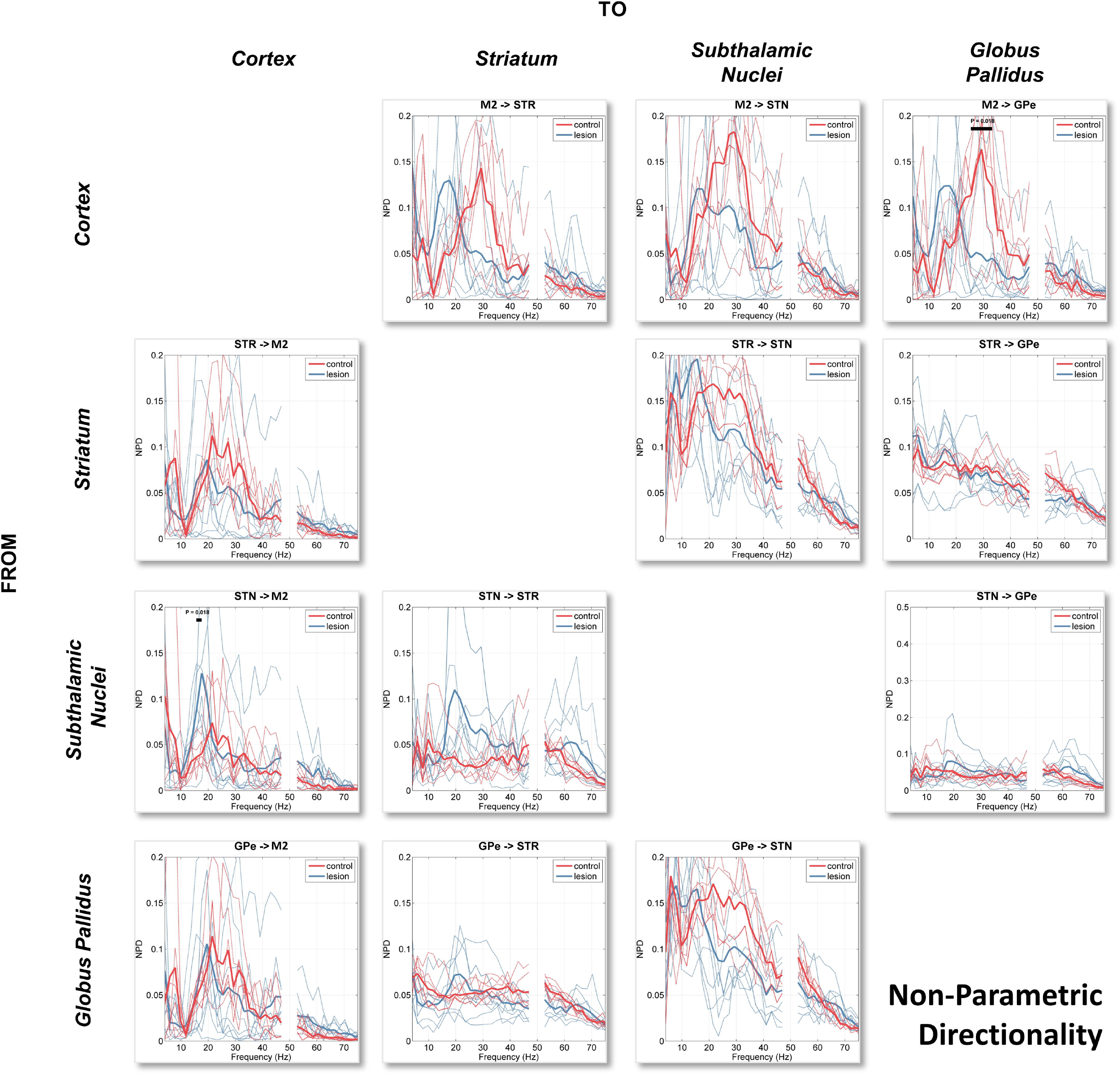
Directed connectivity estimated using non-parametric directionality (NPD) between subcortically recorded LFPs (GPe, STN, and STR) and ECoG recorded from the motor cortex (M2). NPD decomposes the coherence between pairs of signals into forward and reverse components. The array of spectra in the figures reads such that each row title gives the structure with a forward coherence targeted to the structure given by the name given above the column. Spectra from individual animals are shown by the dashed lines and the group means in bold. Different spectra for the control and 6-OHDA lesioned animals are indicated by the red and blue lines respectively.

We observed that directional interactions of low beta-band activity are predominant in the direction leading from M2 and descending the hierarchy of the BG. This effect is most clear in the dopamine-depleted animals. Interestingly we noted a significant difference in the cortical-subthalamic beta band interaction between lesioned and control animals only in the feedback connection STN → M2 (16-18 Hz, P=0.018) which would suggest that STN feedback to M2 is strengthened in the dopamine-depleted state. In the case of the STN/GPe circuit, and unlike iCOH, the non-instantaneous components of NPD do not show 6-OHDA related increases in beta coupling in either direction for the lesioned rats. Rather, NPD suggests a directional asymmetry in activity in the high beta/gamma band with forward connections from GPe → STN connection far stronger than in the reverse direction (cluster statistics testing differences between forward and backward spectra in the 6-OHDA recordings: 4-43 Hz, P<0.001). Notably we see a feedback in the STN → STR that is only present in the lesion condition, a feature that will be relevant with respect to results later in this article.

The pattern of activity in the high beta/gamma range between cortical and subcortical regions appeared to be principally cortically leading with interactions in the 20-40 Hz range being most prominent in the dopamine-intact control rats (top row of figure 5). Cluster-based permutation analysis shows a significant increase in the high/gamma M2 → GPe NPD in the control vs the lesion condition (25-30 Hz, P=0.017). High beta/gamma connections from subcortical structures feeding back to M2, are weaker than the forward connections, but are still present for striatal and pallidal feedbacks to M2 (first column, row 2 and 4, figure 5). Again, there is a clear peak in the high beta NPD from STN to STR in the lesioned animals although a dependence on dopamine was not seen to be significant when cluster statistics were applied. Finding of NPD from STN to STR does not accord the canonical circuit (Figure 1) but may instead imply feedback to striatum via subcortical thalamo-striatal loops that will be discussed later.

### Inferring Routing of Brain Rhythms: Partialized Non-Parametric Directionality

We repeated the NPD analysis as before but this time by systematically partialising out each structure by conditioning the analysis on the LFPs or ECoG recorded at other regions in the network. We again employed cluster statistics to determine significant differences between the non-conditioned NPD spectra (presented in the previous section) and the conditioned variant shown in this section of the results.

#### Conditioning the NPD using Local Field Potentials Recorded from the STN

We first conducted a partialization of the NPD estimate using LFPs recorded from within the STN (figure 6). Conditioning with signals from the STN does not remove beta connectivity between the remaining structures in the network although it does weaken the majority of comparisons in the control (6 of 6 comparisons) but not the lesion (2 of 6 comparisons) experiments (see figure 6, red and blue bars respectively). Cluster statistics indicate that the following NPDs for the control experiments were significantly reduced by conditioning with the STN signal: M2 → STR (14-33 Hz, P<0.001), M2 → GPe (14-33 Hz, P<0.001; 37-49 Hz, P=0.010), STR → GPe (10-49 Hz, P<0.001), GPe → STR (18-49 Hz, P<0.001) as well as feedback connections (returning to cortex): STR → M2 (14-27 Hz, P<0.001), GPe → M2 (18-49 Hz, P<0.001). Furthermore, conditioning the NPD with the signal from STN does not disrupt the 6-OHDA associated increases of M2 input to either the STR (14-21 Hz, P<0.001) or GPe (14-21 Hz, P<0.001). We also found in the dopamine-depleted state that there was increased (relative to the controls) feedback to M2 from both GPe (16-20 Hz, P=0.013) and STR (16-20 Hz, P=0.006).

**Figure 6.**
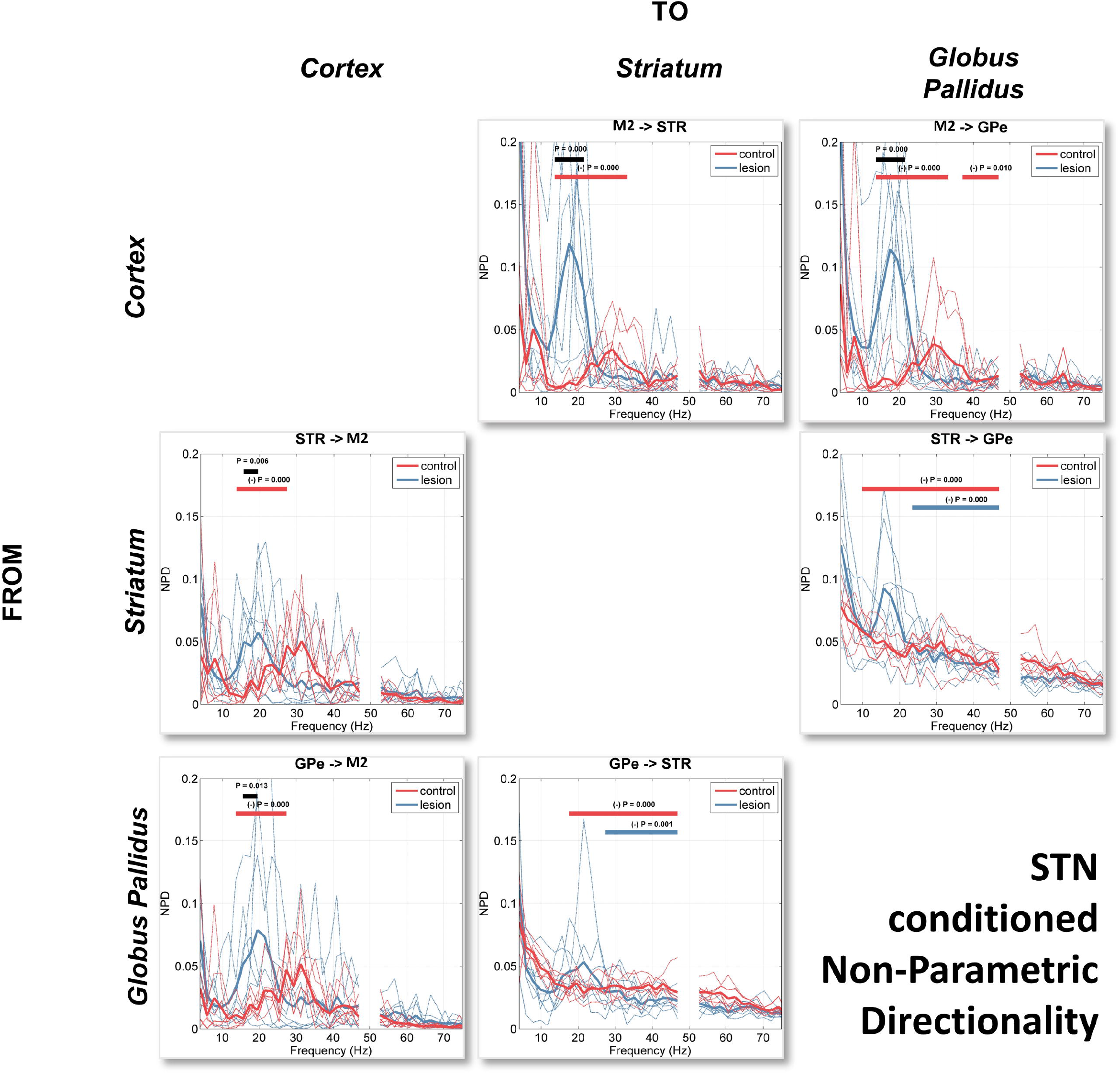
Non-parametric directionality conditioned on LFPs recorded from the STN. Spectra for each animal are shown by thin lines corresponding to either 6-OHDA lesioned (blue) or control (red). The group averages are shown by the bold lines. Cluster-based permutation statistics were applied to determine significant differences between control and lesioned conditions, with significant clusters aindicated by the black bold line above the spectra and corresponding P-value. Significant changes between the original NPD and the conditioned variant shown here are indicated by the coloured bold line corresponding to a difference either with the control (red) or lesioned (blue) signals.

Notably we observed some separation in the effects of the conditioning between the control and lesion experiments. In the control animals conditioning the NPD on LFPs recorded at STN acted to reduce activity in a wide band (~12-40 Hz) for the forward connections (flowing down the indirect pathway; see figure 1), whilst the return connections (STR → M2, and GPe → M2) were only affected by conditioning a tighter band corresponding to low beta. Lesioned animals only showed reductions at higher frequencies (~24-45 Hz, high beta/low gamma) and only between GPe and STR. We observed that conditioning of the NPD with the STN signal acted to significantly reduce interactions between STR and GPe in both the forward (23-49 Hz, P<0.001) and reverse (27-49 Hz, P=0.001) directions.

#### Conditioning the NPD using Local Field Potentials Recorded from the GPe

Next, we performed the NPD analysis of recorded signals but this time conditioning the interactions with LFPs recorded from within the GPe (figure 7). We found that the conditioning had the effect of reducing NPD estimates in 6 out of 6 possible connections in the controls and 3 out of 6 in the 6-OHDA-lesioned rats. Most notably we found that the conditioning significantly attenuated (when compared to the unconditioned NPD) the low beta band interaction in the M2 → STR connection for both recordings made in control (14-39 Hz, P<0.001) and lesioned (14-21 Hz, P<0.001) animals. Secondly, we found a reduction of interactions between STR → STN across a wide range of frequencies, again for both control (6-49 Hz, P<0.001) and lesioned (4-49 Hz, P<0.001) recordings suggesting signal routing is strongly mediated by GPe in accordance with the canonical indirect pathway. Interestingly we found that although beta NPD in the M2 → STN connection was attenuated by conditioning in the control recordings, for the 6-OHDA recordings the prominent low beta peak in the NPD remained and no significant effect of conditioning was observed. Similarly, the STN → M2 feedback also retained a sharp beta peak that remained significantly increased in recordings corresponding to the 6-OHDA lesion experiments (14-20 Hz, P=0.002). Additionally, we found that when conditioning the NPD with the GPe signal, we found that the STR→ M2 demonstrated a significant increase in strength in the 6-OHDA animals (16-21 Hz, P<0.001)

**Figure 7.**
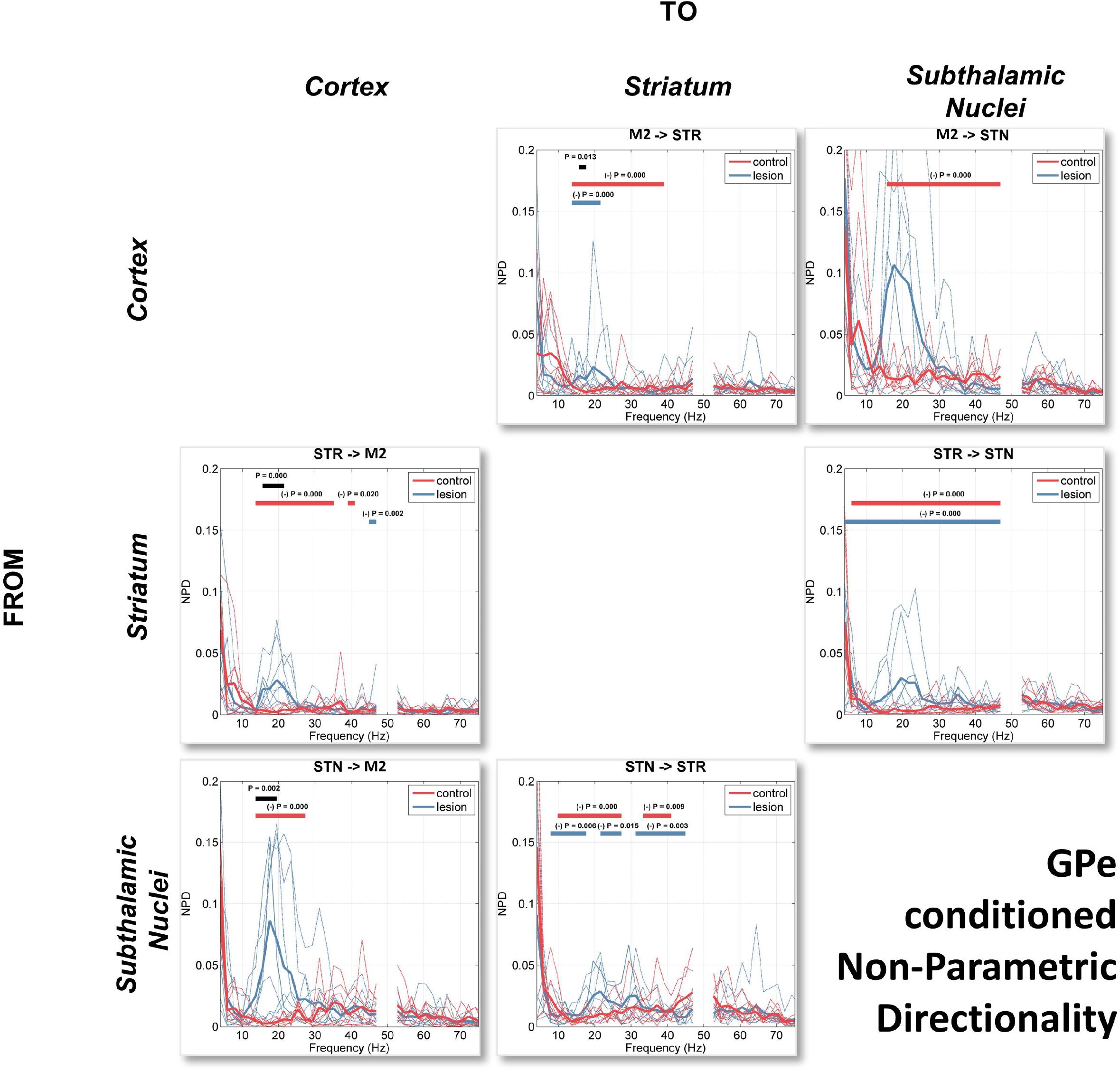
Non-parametric directionality conditioned on LFPs recorded from the GPe. Spectra for each animal are shown by thin lines corresponding to either 6-OHDA lesioned (blue) or control (red). The group averages are shown by the bold lines. Cluster-based permutation statistics were applied to determine significant differences between control and lesioned conditions, with significant clusters aindicated by the black bold line above the spectra and corresponding P-value. Significant changes between the original NPD and the conditioned variant shown here are indicated by the coloured bold line corresponding to a difference either with the control (red) or lesioned (blue) signals.

In the high beta/gamma band we found that conditioning with GPe has a large effect in attenuating the NPD in the forward connections (from M2 descending the indirect pathway) in the control animals: M2 → STR (14-39 Hz, P<0.001), M2 → STN (16-49 Hz, P<0.001), and STR → STN (6-49 Hz, P<0.001). In the lesion animals only, 2 of the 6 comparisons made with NPD were significantly attenuated in the 20-50 Hz range: STR → STN (4-49 Hz, P<0.001) and STN → STR (21-27 Hz, P=0.015; 31-45 Hz, P=0.003). This would imply that interactions in both directions between STN and STR are mediated via GPe.

#### Conditioning the NPD using Local Field Potentials Recorded from the STR

The third set of analyses used the local field potentials recorded at the STR to condition the NPD estimates (figure 8). We found that this had the effect of destroying large parts of the descending interactions (connections from M2 descending the hierarchy of the indirect pathway) in the control animals, namely for M2 → GPe (16-37 Hz, P<0.001) and M2 → STN (16-37 Hz, P<0.001). In the lesion recordings, the effect of conditioning split into two ways: 1) The interactions between the STN/GPe were significantly reduced across a very wide band ranging low-beta to gamma frequencies in both the STN → GPe (8-49 Hz, P<0.001) and GPe → STN (6-49 Hz, P<0.001) condition. 2) Interaction in the “hyper-direct” M2 → STN connection was not attenuated, although the M2 → GPe (likely routed at least in part via the indirect pathway) was suppressed by conditioning with the striatal signal (18-24 Hz, P=0.001). This peak is also seen in the feedback connection from STN → M2 where the significant 6-OHDA associated increase in beta feedback reported in previously analysis was found to remain (18-20 Hz, P=0.012).

**Figure 8.**
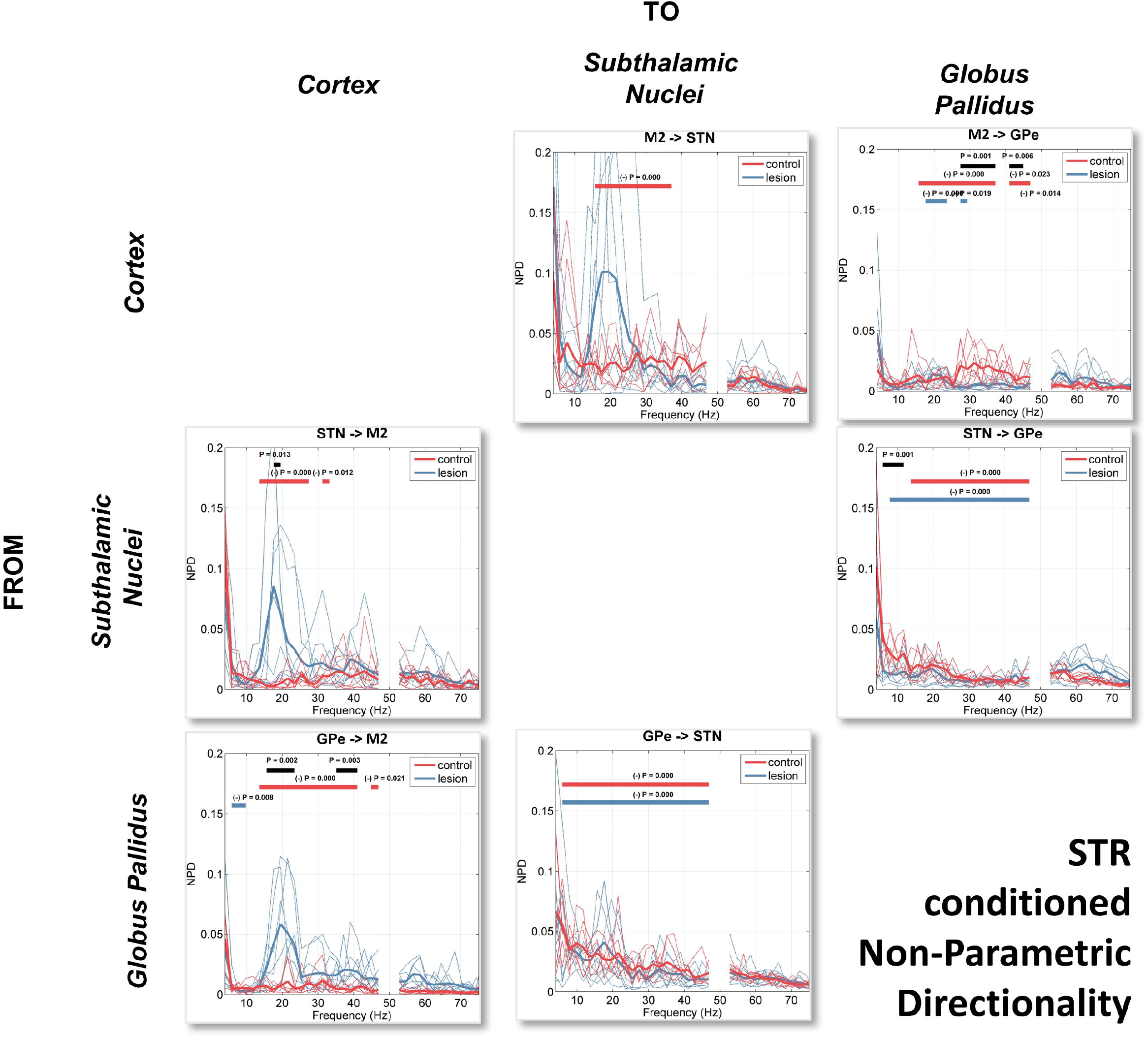
Non-parametric directionality conditioned on LFPs recorded from the STR. Spectra for each animal are shown by thin lines corresponding to either 6-OHDA lesioned (blue) or control (red). The group averages are shown by the bold lines. Cluster-based permutation statistics were applied to determine significant differences between control and lesioned conditions, with significant clusters aindicated by the black bold line above the spectra and corresponding P-value. Significant changes between the original NPD and the conditioned variant shown here are indicated by the coloured bold line corresponding to a difference either with the control (red) or lesioned (blue) signals.

Similar to the NPD estimates conditioned with signals recorded at GPe, we found that the conditioning with LFPs recorded at STR acts to largely remove the high beta/gamma interactions. In the M2 → GPe connection we found that high beta gamma activity was attenuated by conditioning in the control condition (16-37 Hz, P<0.001) as well as a significant condition dependent 6-OHDA associated suppression of activity in this band was observed (27-37 Hz, P=0.001; 41-45 Hz, P=0.005). Additionally, we also found that feedback in this frequency range (for control recordings) from GPe → M2 was significantly attenuated by conditioning with striatal signals (14-41 Hz, P<0.001; 45-49

Hz, P=0.022). Furthermore, we found that activity in this feedback connection was significantly increased in the controls when compared to the 6-OHDA lesioned animals (35-41 Hz, P=0.002).

#### Conditioning NPD Using Field Potentials Recorded from M2

The final analyses utilized ECoG signals recorded from the M2 to condition the NPD estimates (results in figure 9). We found that the NPD estimates conditioned on M2 are greatly flattened and devoid of distinct peaks at either low beta or high beta/gamma frequencies that were seen typically in the other analyses. Altogether 5 of 6 NPD spectra had no distinct spectral peaks. When testing for significant attenuation of NPD following conditioning we found that only control recordings were found to be significantly attenuated. However, the loss of features found in the unconditioned NPD (such as beta or gamma peaks) were equivalent for both the control and 6-OHDA recordings.

**Figure 9.**
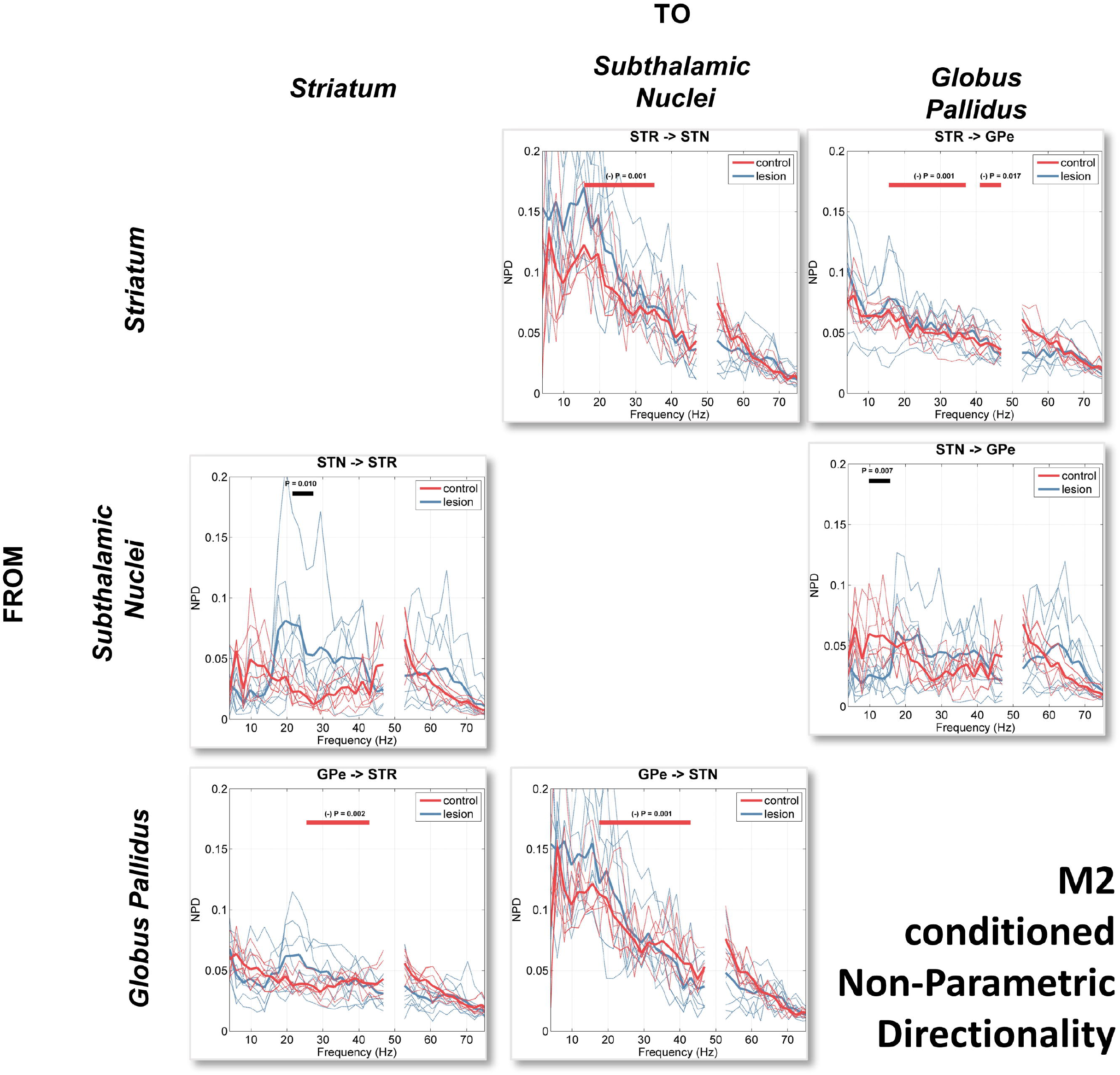
Non-parametric directionality conditioned on the M2 ECoG. Spectra for each animal are shown by thin lines corresponding to either 6-OHDA lesioned (blue) or control (red). The group averages are shown by the bold lines. Cluster-based permutation statistics were applied to determine significant differences between control and lesioned conditions, with significant clusters aindicated by the black bold line above the spectra and corresponding P-value. Significant changes between the original NPD and the conditioned variant shown here are indicated by the coloured bold line corresponding to a difference either with the control (red) or lesioned (blue) signals.

When testing for the effects of 6-OHDA, we found that the STN → STR connection was significantly altered. We observed a broad peak from 20-40 Hz in the lesion recordings that was not attenuated by M2 conditioning and demonstrated a significant increase in strength associated with dopamine depletion (21-27 Hz, P=0.010).

### Summary of Connectivity Analyses

Using recordings made in control and lesioned rats, we identified functional connectivity between cortical and BG sites that involved either low beta or high beta/gamma oscillations. Broadly speaking, we found that gamma connectivity is sensitive to the conditioning of structures upstream of the STN, particularly GPe and STR, which removes any detectable gamma from the spectra. In contrast, beta connectivity is robust to partializing using LFPs of any single structure. Cortico-subthalamic connectivity in the beta range is unaffected by partialising of GPe or STR, suggesting M2/STN low beta connectivity is not routed via the indirect pathway. In the next section, we will outline several putative models of oscillatory dynamics and present evidence from our analyses that either support or weaken the plausibility of each model.

## Discussion

### Hypotheses and evaluation of evidence for signal propagation in the network

We have undertaken a systematic analysis of a dataset involving multisite LFP recordings of the cortico-basal ganglia circuit that contains data from a set of dopamine-intact control rats and another set of rats with chronic dopamine depletion induced by a unilateral injection of 6-OHDA. We will next discuss evidence for competing theories of the propagation of oscillatory activity across the Parkinsonian cortico-basal ganglia circuit.

### Mechanisms of the Flow of Beta Rhythms in the Basal Ganglia Circuit

Here we will evaluate the evidence provided by the analyses reported here in light of a number of proposed theories concerning the generation and propagation of beta-band activity in the network and the changes that occur during dopamine depletion that lead to its amplification. This work is summarised in table 1.

**Table 1.**
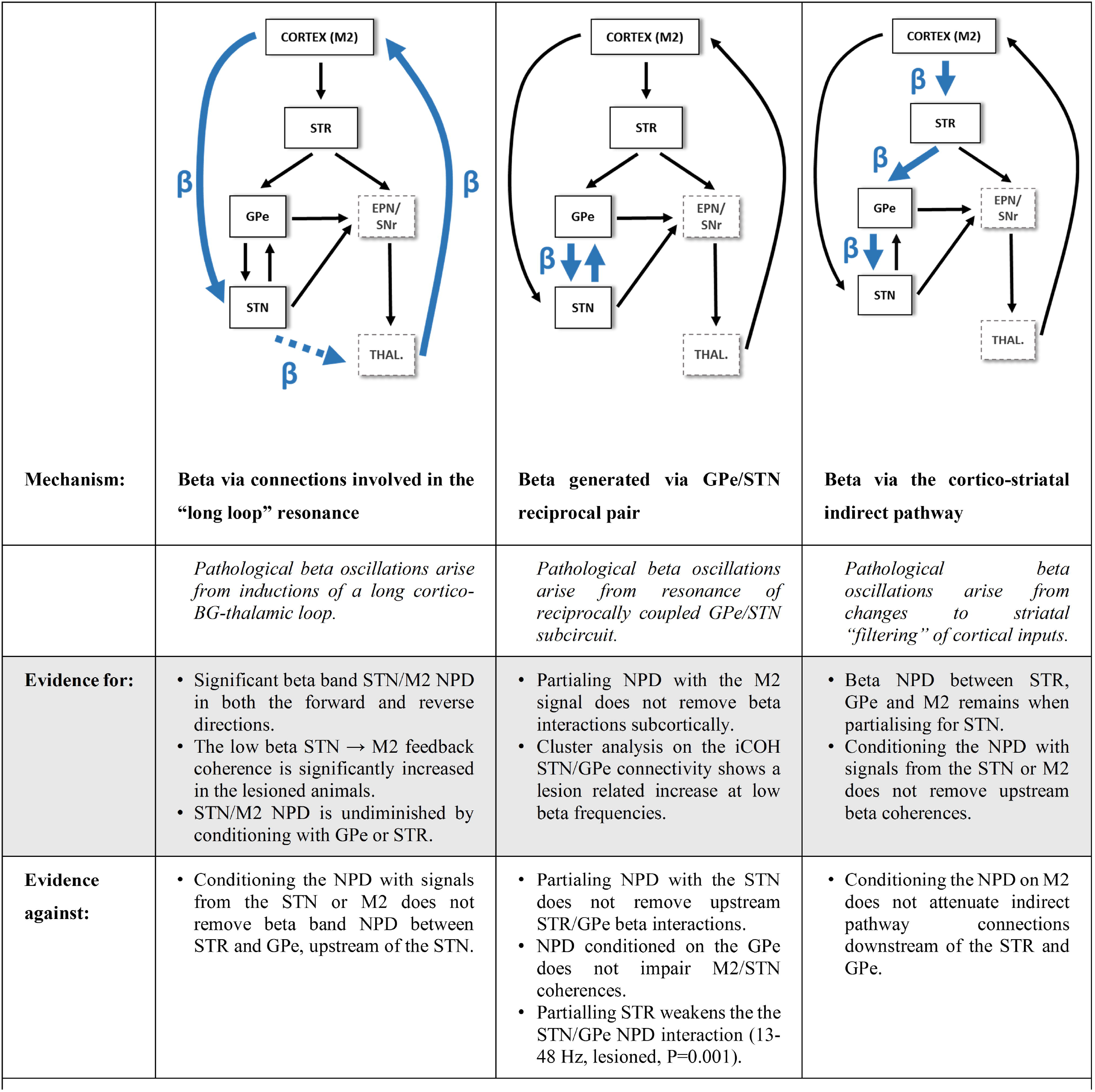
Summary of hypotheses of the impact of dopamine depletion on the propagation of beta rhythms in the cortico-basal ganglia circuit.

#### Hypothesis 1: Dopamine depletion in the basal ganglia induces increased beta resonance in the cortical/STN “long-loop”

Previous authors have suggested that pathological beta rhythms are generated from the strengthening of a long cortical feedback loop that returns from basal ganglia output nuclei via the thalamus. Strengthened coupling is proposed to facilitate pathological resonance at beta frequencies (Brown 2007; van Albada and Robinson 2009; Dovzhenok and Rubchinsky 2012; Pavlides et al. 2015). The first step towards verifying the plausibility of this hypothesis involves determining whether there is indeed functional connectivity between STN and M2 in the beta band, and whether this occurs independently of the cortico-striatal inputs to the indirect pathway.

Analysis of the iCOH for the M2/STN pairing suggests that functional connectivity in the beta band is significantly strengthened in the lesioned animals compared to controls (figure 4). Analysis with NPD has demonstrated that there is a beta peak in the directed coherence in the low beta range in the forward M2 → STN connection for both the control and 6-OHDA animals. However, in the lesioned animals, the feedback connection (STN → M2) is significantly strengthened over that measured in the controls. Neither the hyper-direct M2 → STN connection, nor its feedback (STN → M2) are diminished by either conditioning with signals from the GPe or STR in the lesioned animals (figure 7 and figure 8). This suggests that outflow from STN is routed independently of STR or GPe, likely directly via the BG output nuclei. In contrast, in control rats, NPD of the feedback connections at beta frequencies are significantly decreased by conditioning with the STR signal in the forward (M2 → STN), and backward (STN → M2) directions, suggesting that in the dopamine-intact anaesthetised state, any residual beta band activity is routed via STR, whilst the hyper-direct pathway is relatively quiescent. These findings support the idea that the dopamine-depleted state is associated with a strengthening of the hyper-direct pathway and its subsequent feedback from STN to cortex.

We also provide evidence of multiple coexisting pathways for propagation of beta oscillations. Notably, it was found that conditioning of the NPD with LFPs recorded from the STN (figure 6) does not act to remove the 6-OHDA lesion associated beta NPD in the structures ‘upstream’ of the STN (i.e. the STR and GPe). NPD in the low beta range is significant in both the forward and backward directions along parts of the network involving either M2, STR or GPe. Most notably, we find that striatal-subthalamic interactions are strongly modulated by the GPe signal, a finding in line with the canonical indirect pathway. Future work to validate the long-loop hypothesis would involve the conditioning of the STN → M2 NPD using signals recorded from BG output nuclei (either internal globus pallidus (GPi /EPN in rat) and/or SNr) or their major targets in the thalamus. If these signals were available then it would be possible to determine if there is enhanced cortical return of BG beta rhythms in the dopamine-depleted state.

#### Hypothesis 2: Pathological beta is generated from strengthening of the reciprocally coupled STN/GPe circuit

A separate hypothesis concerning the generation of pathological beta rhythms in the basal ganglia considers the reciprocally coupled STN/GPe circuit from which increased coupling associated with the loss of dopamine induces a pathological beta resonance that spreads across the rest of the network (Plenz and Kital 1999; Bevan et al. 2002; Holgado et al. 2010; Tachibana et al. 2011).

We note that conditioning the NPD with the M2 signal does not remove the strong STN → GPe directed connectivity, but it does attenuate the GPe → STN (figure 9). This would suggest that activity feeding back onto GPe from STN has sufficiently unique information to not be partialized out by the cortical ECoG, perhaps generated by some resonance phenomena. However, a number of the analyses presented here suggest that pathological beta does not originate from STN/GPe circuit resonance. This is summarised as follows: i) Comparison of forward and backward NPD for STN/GPe interactions shows strong asymmetry, with the GPe→ STN connection predominating; ii) conditioning of the NPD using the LFPs recorded at the STR significantly reduces the strength of both GPe → STN and STN → GPe NPDs in both the control and lesion animals (figure 8), suggesting that beta in these structures results from beta propagating through striatum; iii) conditioning the NPD with LFPs recorded at the STN (figure 6) does not act to remove the upstream 6-OHDA associated beta NPD between STR or GPe (although it does significantly weaken beta NPD in the control animals); iv) GPe conditioned NPD analysis does not show impairment of the pathological M2/STN beta interactions (figure 7) suggesting that the beta found at STN can be, at least in part, generated independently of a signal found at GPe. The evidence summarised in point (i), may speak to the hypothesis that the STN/GPe circuit acts as a pacemaker that is in turn tuned by abnormal striatal drive, but by some nonlinear transformation that is not discernible using the conditioned NPD. Taken together we argue these findings give evidence against pathological beta rhythms arising from the pacemaker-like activity of the STN/GPe sub-circuit.

#### Hypothesis 3: Beta arises through aberrant striatal activity and facilitation of downstream hypersynchrony

It has been proposed that aberrant cortico-striatal activity is involved in the emergence of pathological beta rhythms (Kumar et al. 2011; McCarthy et al. 2011; Damodaran et al. 2015). We provide evidence that elements of the BG downstream of STR require a signal that is also shared with the cerebral cortex (M2). In the lesioned animals, there is significant beta band NPD in the M2 → STR and STR

→ GPe connections (figure 5) suggesting a propagation of beta activity down the indirect pathway. With conditioning, we find that the beta peak in the STR → GPe connection is removed in its entirety when conditioning the NPD with the cortical signal (figure 9) in both the control and lesion recordings (interestingly however, the STR → STN beta NPD does remain) suggesting that beta band correlations are at least in part explained by cortical activity. Importantly, we found that conditioning the STR → STN connection with the LFP recorded from GPe (figure 7) acts to significantly reduce the NPD in both control and lesion experiments. This would suggest that beta flow is mediated via GPe in agreement with the canonical view of the indirect pathway demonstrating the ability of partial NPD to discern hierarchical connectivity. Further to this, the finding that upstream GPe/STR beta NPD is susceptible to conditioning with LFPs from STN (figure 6) in dopamine-intact animals may reinforce the idea that in the healthy network, routing of beta occurs predominantly through the indirect pathway, whilst in the pathological case we see beta that is persistent and with activity that is at least in part independent of that recorded at the STN.

#### Hypotheses of the Origins/Routing of High Beta/Gamma Oscillations

The presence of high beta/gamma oscillations in the subcortical network has been noted by a number of authors (Brown et al. 2002; Berke 2009; Sharott et al. 2009; van der Meer et al. 2010; Nicolás et al. 2011) but our understanding of the functional propagation of high beta/gamma oscillations through the network is limited. An evaluation of the evidence is summarised in table 2. We report gamma activity in the LFPs as well as connectivity in the range 30-60 Hz which is in good agreement with that previously reported in anaesthetised rats (Magill et al. 2004; Sharott et al. 2005b, 2009). Activity in the awake rats, during movement has also been reported, albeit at slightly higher frequencies (Brown et al. 2002; Brazhnik et al. 2012; Delaville et al. 2014).

**Table 2.**
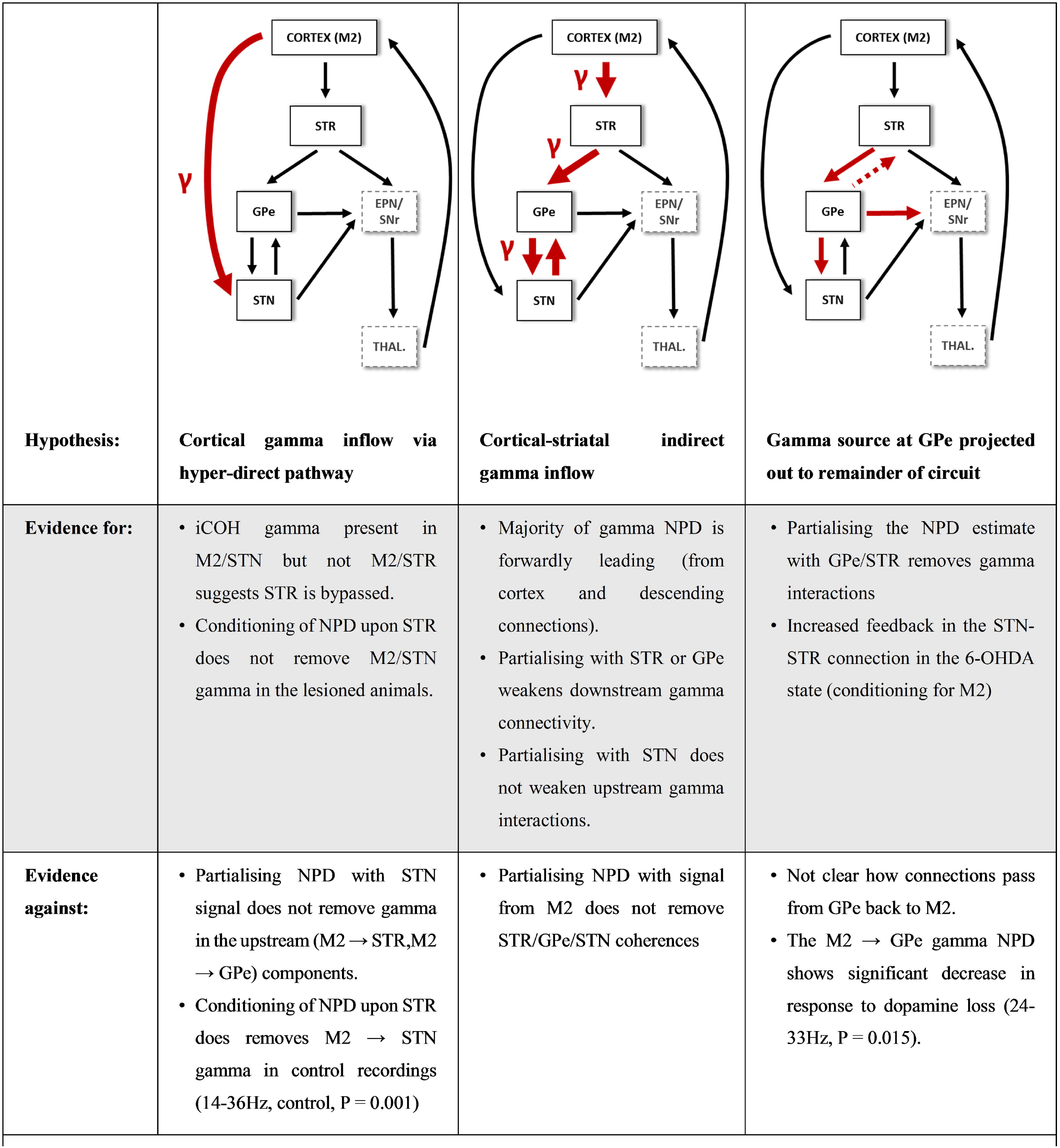
Summary of hypotheses for gamma flow in the cortico-basal ganglia circuit.

#### Hypothesis 4: High beta/gamma enters the subcortical network via the hyper-direct M2 → STN connection

Results from analyses which used iCOH to investigate non-zero lag correlations between BG structures and the cortex suggested that gamma interactions are routed in a way that bypasses STR as a gamma peak is absent in the M2 (figure 4). The hyper-direct pathway is the other principal source of cortical input to the BG, therefore the marked weakness of gamma interaction in the M2/STR when compared to the M2/STN iCOH spectra may imply that the hyper-direct pathway is responsible for gamma input to the network.

However, whilst there is a large peak in the high-beta/low-gamma band NPD for the M2 → STN interaction(figure 5), if we examine the same connection but conditioned on LFPs either recorded at STR (figure 7) or GPe (figure 8) we see that the conditioning significantly reduces NPD in the control animals (M2 → STN conditioned on STR-16-37 Hz, P<0.001; M2 → STN conditioned on GPe 16-47 Hz, P<0.001) suggesting any directed coherence between M2 and STN in these animals is routed via striatal-pallidal connections. Furthermore, if we condition the NPD with LFPs recorded at the STN (figure 6), we see that gamma interactions remain in the upstream components (M2 → STR, M2 → GPe) again suggesting striatal-pallidal connectivity is vital in the propagation of gamma rhythms. When taken together, these data do not supply strong evidence that the source of high beta/gamma input in the network is supplied by a hyper-direct cortico-subthalamic route.

#### Hypothesis 5: Gamma enters the network via cortico-striatal inputs and reaches STN via the indirect pathway in a dopamine dependent manner

An alternative to high beta/gamma oscillations entering via hyper-direct STN input is that they are channelled via the cortico-striatal indirect pathway. The clearest results of the NPD analysis in the high beta/gamma band can be seen to be for the forward NPDs originating from M2 and passing on to the subcortical regions (figure 5). Connections M2 → STR, M2 → GPe, and M2 → STN all show high values of NPD in this frequency band (> 0.15) suggesting that most of the gamma is directed from the cortex. Furthermore, conditioning the NPD with either LFPs recorded at the STR (figure 8) or GPe (figure 7) acts to remove gamma interactions both upstream and downstream of the STR (with respect to the indirect pathway). Subsequently, conditioning of the NPD with STN (figure 6) was less effective at attenuating gamma band interactions than when using signals higher in the indirect pathway, suggesting that the gamma descends the hierarchy, from either a cortical or striatal source. Notably, we observed that STN conditioned NPD did not act to attenuate feedback connections from GPe or STR back to the M2. This would suggest routing of gamma to the M2 in a way that occurs independently of STN.

In attempt to elucidate the source of the gamma activity we conditioned the NPD on the cortical ECoG (figure 9). We find that gamma connectivity in the control recordings, dopamine depletion acts to significantly reduce NPD coefficients for the GPe → STN and STR → STN connections, yet the feedback connection STN → STR is unaffected. This connection in the control animals shows a peak from 18-42 Hz which is significantly larger than in the lesioned animals. This is in agreement with the idea of prokinetic gamma and in accordance with findings in patients (Sharott et al. 2014). Furthermore, these findings suggest that gamma activity is directed to upstream components of the BG in a way independent of M2, perhaps via a subcortical feedback loop.

#### Hypothesis 6: High beta/gamma is generated in the GPe or STR and downstream to STN

The finding that conditioning the NPD with cortical ECoG does not entirely abolish gamma connectivity within the BG may suggest that at least some high beta/gamma propagates locally within the BG. When conditioning was performed with LFPs recorded from the GPe (figure 7) or STR (figure 8), we found that interactions in the high beta/gamma frequency ranges were abolished for the majority of subcortical interactions. This would imply that these GPe and STR structures are necessary for the propagation of high beta/gamma interactions in the both the control and 6-OHDA lesion animals. This in combination with the evidence provided for hypothesis 5 suggests that high beta/gamma originates at either STR or GPe and then propagates to downstream structures. Backward gamma interactions from GPe to STR are apparent in the NPD conditioned on M2 or STN, suggesting the STR signal is the result of local propagation of a gamma signal from GPe. From the canonical view of the circuit it is not clear how gamma passes upstream from GPe. However, a substantial proportion of GPe neurons that innervate the striatum have been shown to exist, with one GPe cell type (arkypallidal neurons) projecting exclusively to striatum (Mallet et al., 2012; Abdi et al., 2015; Hegeman et al. 2016).

### Summary of Findings

In this paper, we have presented analyses that investigate propagation of oscillatory activity through connected regions of the cortico-basal ganglia network. We have applied a novel method of partialized directed coherence to achieve a systematic deconstruction of the propagation of rhythmic activity between regions of the network inferred from the LFPs and ECoGs recorded at multiple sites within that network. Using the 6-OHDA-lesioned rat model of Parkinsonism, we demonstrate marked differences in the functional connectivity that result as a consequence of dopamine depletion.

We find diffuse beta synchronization of LFPs across the network that is strongly associated with chronic dopamine depletion. With regards to functional beta connectivity in the network we find evidence for:

1. An increased cortical drive to the basal ganglia following dopamine depletion.
2. Significant beta-band connectivity between structures interacting with the STN that is independent of activity upstream in the indirect pathway (at STR and GPe). This likely originates from the ‘hyperdirect’ cortico-subthalamic input.
3. Increase in feedback of BG structures to M2 after dopamine depletion, proffering evidence in favour of a hypothesis of dopamine-dependent modulation of the long re-entrant cortico-BG-thalamo-cortical loop.
4. Activity dynamics of the STN/GPe subcircuit being partly dependent upon drive from striatum.
5. A feedback from STN to STR that is independent of M2 and significantly strengthened after dopamine depletion, suggesting a strengthening of recurrent subcortical circuits.

Furthermore, we provide evidence for the existence of high beta/gamma synchrony within the network, with evidence that dopamine depletion acts to weaken these rhythms. We summarise our findings with respect to high beta/gamma band interactions in the following:

1. Gamma propagates down the indirect pathway from STR to GPe to STN. This activity is likely generated at cortex.
2. At least some of the gamma activity found at STN is independent of M2 and provides evidence for a subcortical return of subthalamic outputs back to striatum.
3. At least some of the gamma activity returning to cortex is independent of STN, perhaps indicating propagation through the direct pathway.

#### Propagation of Low Beta via two Coexisting but Distinct Streams

In the case of low beta oscillations, we find our data most strongly support a hypothesis that in the dopamine-depleted condition, beta propagation in the network is biased to favour low beta synchrony via induction of a long cortico-subthalamic loop that inputs to the BG via the hyper-direct pathway. Furthermore, we see strong evidence that the return connection from STN to M2 is significantly stronger in the lesioned animals than in the dopamine-intact controls. This provides supporting evidence for the notion that pathological beta amplification arises from entrainment of the re-entrant cortical/STN loop (Brittain and Brown 2014). We speculate that strengthening of the hyper-direct input acts to “short-circuit” the network, such that transmission of information along the indirect pathway is compromised. In the “hold your horses” model of the STN’s role in decision making (Frank 2006; Frank et al. 2007), the hyper-direct pathway is proposed to provide a cortical veto signal which may act to suppress premature action. In the case of PD, over activity of this circuit via increased resonance may act to lock the network into a state that ultimately supresses action. These findings are in agreement with previous research which have found good evidence for bidirectional connectivity between STN and cortex (Lalo et al. 2008; Jávor-Duray et al. 2015).

This hypothesis requires further testing through analysis of the role of the BG output nuclei at GPi or SNr (or their targets in the thalamus) in the propagation of activity. This could be achieved using a functional ‘lesion’ approach like that described in this paper. Furthermore, biophysical modelling of the cortico-subthalamic loop may yield insight as to whether this is plausible given the known conduction delays for the connections in the network. Long feedback loops involving cortex have been demonstrated to be capable of generating oscillatory activity (Leblois et al. 2006; Pavlides et al. 2015). However, work by Shouno et al. (2017) suggests that the required delay for the return of the beta oscillation from STN to cortex may be too large to support resonance in the low beta band. Perhaps the engagement of shorter subcortical loops either subcortical-thalamic loops (McHaffie et al. 2005); or activity of recurrent subthalamo-striatal projections (Sato et al. 2000; Koshimizu et al. 2013) may be more suitable candidates.

The analysis presented here also suggests that a cortico-subthalamic pathway is not the exclusive pathway for beta rhythms within the network, yet may be necessary for enhancement of the STN feedback to cortex that may induce pathological resonance. We would suggest that both the hyperdirect and indirect routes for beta propagation coexist. These two pathways could originate form (be driven by) distinct populations of cortical projections neurons (namely those of the pyramidal tract and intra-telencephalic projections, respectively) and so are likely to show a degree of independence from one another. The data presented here also suggest a second pathway upstream of STN involving the STR that is most evident in the recordings from control rats. We suggest that both pathways contain signals shared by activity measured in the cortical ECoG as conditioning of the NPD acts to remove beta peaks from the majority of connections that were analysed, leaving just beta coherence at the STR → STN connection. These findings support the hypothesis that dopamine cell loss acts to increase the sensitivity of the STR to cortical inflow acting to gate beta to the remainder of the circuit (Magill et al. 2001; Tseng et al. 2001).

Notably, our data do not support the hypothesis of beta generation via an autonomous STN/GPe pacemaker network, as directional coherence between the two is heavily attenuated by conditioning with LFPs recorded upstream in the STR and there is significant asymmetry in the NPD with the pallidal drive predominating. In agreement, Moran et al. (2011) found evidence for a weakening of the STN to GPe feedback connection in the dopamine depleted state, conflicting with the STN/GPe resonance hypothesis. Estimates of effective connectivity from DCM studies have also suggested that input from cortex to STN is strengthened in the Parkinsonian state (Moran et al. 2011), a finding in line with the idea that dopamine enforces cortical influence upon the STN/GPe network (Magill et al. 2001; Leblois 2006; Leblois et al. 2006; Holgado et al. 2010). It may be the case that during PD cortical activity subsumes the STR as the primary driver of the STN/GPe subcircuit. It has been demonstrated that movement is associated with a decreased cortico-pallidal coherence during movement in humans (van Wijk et al. 2017) suggesting disengagement of this circuit is pro-kinetic. Thus pathological resonance may arise as a result of alterations to striatal output (Kumar et al. 2011; Damodaran et al. 2015) that in the healthy system act to decorrelate spiking activity between the two structures (Wilson 2013). The action of dopamine upon these inputs is likely to modulate the existence of beta amplifying phase alignments between STN and GPe such as that observed by Cagnan, Duff, & Brown (2015).

#### Dopamine Depletion is Associated with an Increased Subthalamo-Striatal Feedback

Taken together, the analyses presented here speak to the existence of a high beta/low gamma rhythm that is in general reduced by dopamine depletion. Specifically, our results indicate a band at 27-34 Hz is attenuated in the lesioned rats. Experiments investigating LFPs in the motor cortex of moving rats have demonstrated an increase in activity in this band during movement suggesting that activity at these frequencies in the M2 and SNr is prokinetic (Brazhnik et al. 2012). Our data would suggest that high beta/gamma activity in the normal network is predominantly driven by the cortex as evidenced by: i) the unconditioned NPD indicates that gamma is prominently in the forward direction leading from cortex to subcortical sources; and ii) conditioning the NPD on ECoG recorded at M2 acts to diminish the subcortical directional coherence across a wide band for all connections not involving STN. However, studies by Zold and colleagues has demonstrated that oscillatory activity >20 Hz in corticostriatal afferents is not effectively transmitted (Zold et al. 2012).

Furthermore, following partialization some interactions involving STN do remain. In particular we provide evidence for a significant strengthening of feedback from STN to STR in the lesioned animals in the high beta/gamma band. We speculate that this signal is facilitated through the strengthening of subcortical loops such as that of the thalamo-striatal pathways (McHaffie et al. 2005). Thalamic afferents make up to at least 25% of input onto spiny projection neurons in the STR (Doig et al. 2010; Smith et al. 2014) but have been far less studied than cortical inputs. Work investigating synaptic remodelling following 6-OHDA depletion in mice has suggested that thalamo-striatal inputs to medium spiny neurons are shifted in favour of the indirect pathway (Parker et al. 2016) perhaps enhancing striatal return of subthalamic activity in a mechanism independent of cortex.

### Study Limitations

#### Incomplete Signals for Conditioning

The use of partial coherence for inferring neural connectivity is not in itself a novel approach (Rosenberg et al. 1998; Eichler et al. 2003; Salvador et al. 2005; Medkour et al. 2009), and the application of the conditioned NPD to LFPs recorded in the rat hippocampus has been previously reported (Halliday et al. 2016). However these analyses assume that the signals used for conditioning completely capture the activity going through the proposed pathway. This however is unlikely to be completely the case due to the finite sampling of the structures afforded from the use of electrodes. However, the large number of channels for recording ensure that multiple samples are obtained from each structure. In the data presented here, subcortical structures were recorded from between 2-8 different channels which were all used to conditioning the estimate of directed coherence. It must also be noted that this limitation is likely to apply most to the larger structures that were analysed, namely the motor cortex and striatum, whereas recordings from the smaller sized STN is more likely to capture the majority of activity. This must be considered when interpreting conditioning of the NPD with respect to STR signals. It could be the case that M1 → STN connectivity remains as a result of incomplete sampling of striatal signals.

#### Inference of Connectivity from Non-Spiking Brain Activity

This study focuses upon conclusions that are drawn from mesoscale recordings of brain activity as measured either in the ECoG or LFPs. Transmission of information in the brain is facilitated by axonal propagation of action potentials that are not explicitly captured in these signals. LFPs and ECoG comprise a conglomerate of sub- and supra- threshold events that may or may not be tied to spike activity and so direct inference of neurophysiological connectivity *per se* is limited by this. Nonetheless, it has been previously demonstrated by Mallet and colleagues that beta activity in the LFPs recorded at STN and GPe are associated with increased synchronization of action potential firing by neurons in these structures (Mallet et al. 2008a, 2008b). Furthermore, we provide evidence for the existence of temporally lagged correlations between rhythmic local field potentials recorded between distinct regions of the cortico-BG network that imply causation from one signal to another, a phenomenon that would itself not be possible without the transmission of action potentials. Future work will require the investigation to determine whether directional interactions are ascertainable from multiunit activity and how this relates to lagged synchronization of LFPs.

#### Limits to Inference of Directed Connectivity from Neurophysiological Signals Alone

In this paper, we aim to infer how neural activity propagates across the BG network by investigating the statistical relationships between brain signals. The challenges that this approach face are well documented (Friston 2011; Bastos and Schoffelen 2016). With respect to this study, the benefits that that we put forth for using a model free, non-parametric approach (namely agnosticism to the underlying generating mechanisms of the data) may in turn limit the degree of inference that can be made. Estimates of directed functional connectivity in this paper follow from the assumptions that temporal precedence is indicative of causation. It is however well documented that zero lag synchronization can emerge from neural circuits with particular (but not unusual) network motifs (Vicente et al. 2008; Viriyopase et al. 2012; Gollo et al. 2014). Additionally, “anticipatory” synchronization in which positive lags arise from a directed input have also been described in theoretical neural dynamics (Ambika and Amritkar 2009; Ghosh and Roy Chowdhury 2010). The anatomically tightly coupled STN-GPe subcircuit is a prime candidate for which these phenomena may permit vanishingly small phase lags that make the interactions blind to NPD. Answers to these problems may be given in the future by the fitting of biophysical models to the data presented in this paper. This would provide a well-defined, quantitative description of the potential mechanisms that act to generate the phenomena we have described.

## Conclusion

Overall, we provide a systematic deconstruction of the propagation of pathological rhythms across the Parkinsonian cortico-basal ganglia circuit *in vivo*. These findings strengthen our understanding of how normal and pathological rhythms propagate across the network. Our work highlights the importance of considering non-canonical connections in the network, in particular the activity of recurrent subcortical projections that may act to amplify pathological activity within the BG. Future work will aim to understand the exact changes to the network required to generate the patterns of functional connectivity presented here, as well as to investigate the relationship with spiking activity in the network.

## Acknowledgments

We thank Dr N. Mallet for acquiring some primary data sets.

## Funding

Medical Research Council UK (awards UU138197109, MC_UU_12020/5 and MC_UU_12024/2 to P.J.M.; MC_UU_21024/1 to A.S.). Parkinson’s UK (Grant G-0806 to P.J.M.). S.F.F. receives funding from UCLH BRC. Engineering Research Council UK (awards EPSRC EP/F500351/1 to T.O.W.; EP/N007050/1 to D.H.). The Wellcome Trust Centre for Neuroimaging is funded by core funding from the Wellcome Trust (539208).

